# VariCarta: a comprehensive database of harmonized genomic variants found in ASD sequencing studies

**DOI:** 10.1101/608356

**Authors:** Manuel Belmadani, Matthew Jacobson, Nathan Holmes, Minh Phan, Paul Pavlidis, Sanja Rogic

**Affiliations:** Michael Smith Laboratories, UBC, Vancouver, BC; Department of Psychiatry, UBC, Vancouver, BC

## Abstract

**Background:** Recent years has seen a boom in the application of the next-generation sequencing technology to the study of human diseases, including Autism Spectrum Disorder (ASD), where the focus has been on identifying rare, possibly causative genomic variants in ASD individuals. Because of the high genetic heterogeneity of ASD, a large number of subjects is needed to establish evidence for a variant or gene ASD-association, thus aggregating data across cohorts and studies is necessary. However, methodological inconsistencies and subject overlap across studies complicate data aggregation.

**Description:** Here we present VariCarta, a web-based database developed to address these challenges by collecting, reconciling and consistently cataloguing literature-derived genomic variants found in ASD subjects using ongoing semi-manual curation. The careful manual curation combined with a robust data import pipeline rectifies errors, converts variants into a standardized format, identifies and harmonizes cohort overlaps and documents data provenance. The harmonization aspect is especially important since it prevents the potential double-counting of variants which can lead to inflation of gene-based evidence for ASD-association.

**Conclusion:** VariCarta is the largest collection of systematically curated, harmonized and comprehensively annotated literature-derived ASD-associated variants. The database currently contains 35,615 variant events from 8,044 subjects, collected across 50 publications, and reconciles 6,057 variants that have been reported in literature multiple times. VariCarta is freely accessible at http://varicarta.msl.ubc.ca.

## Background

The genetics risk of ASD is based on the interplay between common inherited variants and rare, often *de novo*, variants. While the aggregated contribution of common variants is estimated to be high, ranging up to 50% (Klei et al., 2012; Gaugler et al., 2014), their individual effect sizes are small and only a few common risk loci have yet been identified so far, partially due to the limited cohort sizes used in genome-wide association studies up to now (He et al., 2013a; Grove et al., 2017; Kosmicki et al., 2017; Vorstman et al., 2017). On the other hand, rare variants with large effect sizes, such as copy-number variants or likely gene damaging single nucleotide variants (SNVs), were found in many ASD individuals; it is estimated that rare genetic variants, both *de novo* and inherited, are causal in at least 10–30% of cases (Iossifov et al., 2012, 2014; Sanders et al., 2015; Turner et al., 2017a; Vorstman et al., 2017; Yuen et al., 2017). Consequently, most of the effort towards understanding the etiology of ASD in the last half a decade has gone into employing next-generation sequencing technology to identify rare, possibly causative mutations in ASD individuals.

There have been a few dozen whole exome sequencing (WES) or whole genome sequencing (WGS) studies of various ASD cohorts, reporting on identified variants and genes containing them. The overlap within and between the studies’ findings at the gene and, especially, variant level has been limited, highlighting the genetic heterogeneity of the disorder. Thus a key strategy for identifying ASD candidate genes is to sequence large numbers of individual genomes to identify likely impactful events, especially *de novo* mutations, recurring in the same gene in unrelated individuals (Buxbaum et al., 2012; Sanders et al., 2012a). This approach resulted in the discovery of many ASD candidate genes. Current estimates for the total number of ASD-associated genes range from several hundreds to over a thousand (Iossifov et al., 2012, 2014; Sanders et al., 2012b, 2015; He et al., 2013b; Ronemus et al., 2014).

While the sequencing efforts continue, there is a need to aggregate and consolidate the findings. The analysis and reporting of variants across studies can vary in many respects, such as reference assembly, transcript set, variant calling and annotation software used, as well as the choice of variant nomenclature, and without careful curation and standardization they are not easily comparable and integrable (Deans et al.; McCarthy et al., 2014; Yen et al., 2017). At the same time, some cohorts (or individuals within cohorts) have been repeatedly sequenced and analyzed, and the same variants have been reported multiple times across publications, but not necessarily in a consistent format. If pooled across studies, this can lead to double-counting and inflation of gene-based evidence for ASD association. With these challenges in mind, we developed VariCarta with the goal of collecting, reconciling and accurately cataloguing literature-derived ASD-associated variants. We employ precise, systematic curation of the data, standardized processing and reporting, identification of overlaps, and comprehensive annotation. The harmonized data is available to researchers for querying and download at http://varicarta.msl.ubc.ca.

## Construction and content

We searched the literature for publications reporting rare genomic variants, SNVs and InDels, found in subjects with ASD diagnosis. We prioritized whole-genome and whole-exome studies (37) over candidate gene studies (13). Papers currently included in VariCarta are listed in Supplemental File 1.

For each publication, we applied the following curation procedures, with an intermediate goal of organizing all relevant variant information in a tabular format that is ready for import. The first step in this process is to copy the relevant text from the source file (typically a supplementary file) as-is in a template import document. The completed document is composed of a set of predefined worksheets, which contain the publication’s metadata, variant data and a description of the steps needed to automatically extract, transform and load the data into a uniform variant data model. This document is parsed by a computational pipeline, which validates and stores the data into a relational database. The link to the pipeline’s source code as well as an example import document are found at varicarta.msl.ubc.ca/downloads.

There are different variant reporting conventions used in in the literature, so careful inspection is needed to determine the format that was used in a publication, and steps needed to convert it into a standardized format VariCarta uses for displaying variants. These steps are described in the import document and may include conversion of non-genomic to genomic variant coordinates using TransVar (Zhou et al., 2015), biocommons UTA and HGVS packages (Hart et al., 2015), UCSC liftOver of genomic coordinates to GRCh37/hg19 human genome assembly, conversion from zero-based to one-based indexing system, correction of erroneous reference bases and so on. Once the genomic coordinates have been resolved and the uniform formatting has been applied, we use ANNOVAR (Wang et al., 2010) to annotate variants. This annotation includes gene-level information, such as genomic context, transcript accession number, and functional effect of the variant within the gene. We also include variant-level annotations using CADD (Kircher et al., 2014) for variant deleteriousness prediction and ExAC (Lek et al., 2016) for allele frequencies in the general population. At this stage, we exclude variants that are found in control subjects or subjects not reported to have an ASD diagnosis (for studies that report on cohorts with different neurodevelopmental phenotypes). Variants were excluded if they were reported as having failed to validate using an orthogonal sequencing technology, whose coordinates and/or reference alleles cannot be confidently disambiguated or were problematic for some other reason detailed in the curator notes. Although the trend in ASD sequencing studies is to report mainly rare coding genomic variants, the final list of imported variants may include common and non-coding variants if these were present in the original report. These are easily distinguished based on ExAC frequencies or genomic context.

The last phase of variant processing is harmonization, which ensures that variant events are not double-counted. We define a “variant event” as a unique combination of a reference allele, its genomic location and alternative allele belonging to a single individual. We also define a “complex event” as a grouping of two or more variant events from the same individual that differ but have overlapping or adjacent genomic coordinates. This indicates that the grouped variant events might be describing the same underlying genotype but are incongruent due to the heterogeneity of formats used across papers. For example, the same several-base long substitution can get reported either as a single substitution variant or a series of SNVs.

Because subject IDs are used to define variant events, we take special care when handling cohorts that are already present in VariCarta to ensure that subject IDs between the studies are consistent. Occasionally, the same subject could have slightly different IDs between studies: for example, Yuen et al. 2015 (Yuen et al., 2015) uses “-” (dash), while Yuen et al. 2017 (Yuen et al., 2017) uses “_” (underscore) in otherwise identical subject IDs from the same cohort. Another example is the omission of “p1” suffix, meant to indicate a proband, from a Simons Simplex Collection (SSC) family ID in Iossifov et al. (2012) (Iossifov et al., 2012), Iossifov et al. (2014) (Iossifov et al., 2014) and Ji et al. (2016) (Ji et al., 2016). We attempt to rectify such discrepancies during the curation stage. This is facilitated by the introduction of internal VariCarta-specific subject IDs that can be linked with more than one original study ID. However, we were not always able to resolve overlaps, such as in cases were subject IDs are not provided or are non-mappable to the original cohort identifiers. While these variants appear in the database as uniquely reported, the curation notes on the publication details page, would indicate the possibility of overlap.

## Utility and discussion

### VariCarta web site features

The central feature of the website is its capability of querying variants by gene name/symbol or by genomic region. The resulting variant table shows the summarized information about all the variants found in the query region and is the launching point for accessing gene information from the Ensembl and NCBI databases, genomic annotation of the region using the UCSC Genome browser, information about papers that reported the variants and original, published variant data. A variant event is displayed only once in the variant table, with the Sources column listing IDs of all publications reporting it. The publication IDs are tagged to indicate the scope of sequencing study ([W] whole genome sequencing, [E] whole exome sequencing, [T] targeted sequencing). A complex event is initially displayed as one row that can be expanded to show the information about each grouped variant event (an example of a complex event is shown in Table 1). All query results are available for download as plain text/csv format from the spreadsheet icon in the table navigation header bar.

**Table 1:**
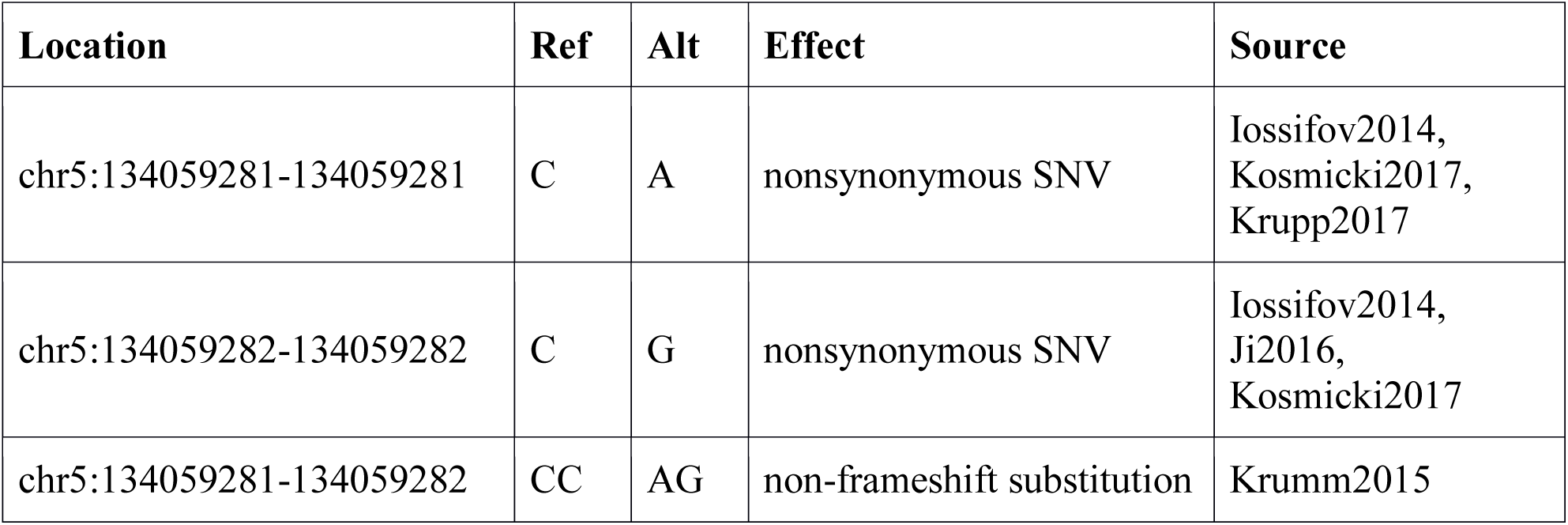
An example of a complex event in VariCarta

A two-base substitution found in SSC subject 14606.p1 has been reported in five different papers included in VariCarta. Iossifov et al. (Iossifov et al., 2014) and Kosmicki et al. (Kosmicki et al., 2017) report it as two contiguous SNVs (C->A and C->G) at positions 134059281 and 134059282 on chr5, while Krumm et al. (Krumm et al., 2015) reports it as a non-frameshift substitution (CC->AG). Both representations describe the same resulting genotype. The other two papers report just one of the contiguous SNVs each (Krupp et al. (Krupp et al., 2017): C->A and Ji et al. (Ji et al., 2016): C->G). This complex variant can also be viewed directly in VariCarta at varicarta.msl.ubc.ca/variant?chr=5&start=134059281&stop=134059281.

The original data as presented in the source publication for each variant event can be accessed by clicking on a magnifying glass icon, which launches a pop-up window displaying the variant information parsed from the publication of origin or inferred during the curation stage. This allows users to confirm the relationship between the source and our representation of it in the VariCarta system.

The list of publications that have been curated and included in VariCarta is shown on the Publications page (varicarta.msl.ubc.ca/publications; also see Supplemental File 1). The basic publication information is listed in a table with links to the original publications. Clicking on a book icon in the Details column opens a page with more detailed information about the study, including methodology used, size and type of the cohort, and types of variants reported. We also provide curation notes, which detail issues that had to be resolved during the curation stage, or other noteworthy information regarding the study. Clicking on the variant event count will display all the variants from that publication that are available in VariCarta.

The Statistics page (varicarta.msl.ubc.ca/stats) offers several gene rankings based on different criteria. These rankings are not intended to be used for a prioritization of ASD candidate genes because they can include common variants (non-coding variant are excluded from ranking calculations), as well as variants from targeted studies, which are typically excluded for the prioritization purposes. The Statistics page also shows the distribution of variants across publications, functional effects and genomic features. Finally, it includes a heatmap of the variant overlap between publications (Figure 1), which illustrates the extent of variant double-reporting across the literature. The variants reported in two different papers can be accessed by clicking on the overlap number in the heatmap.

**Figure 1:**
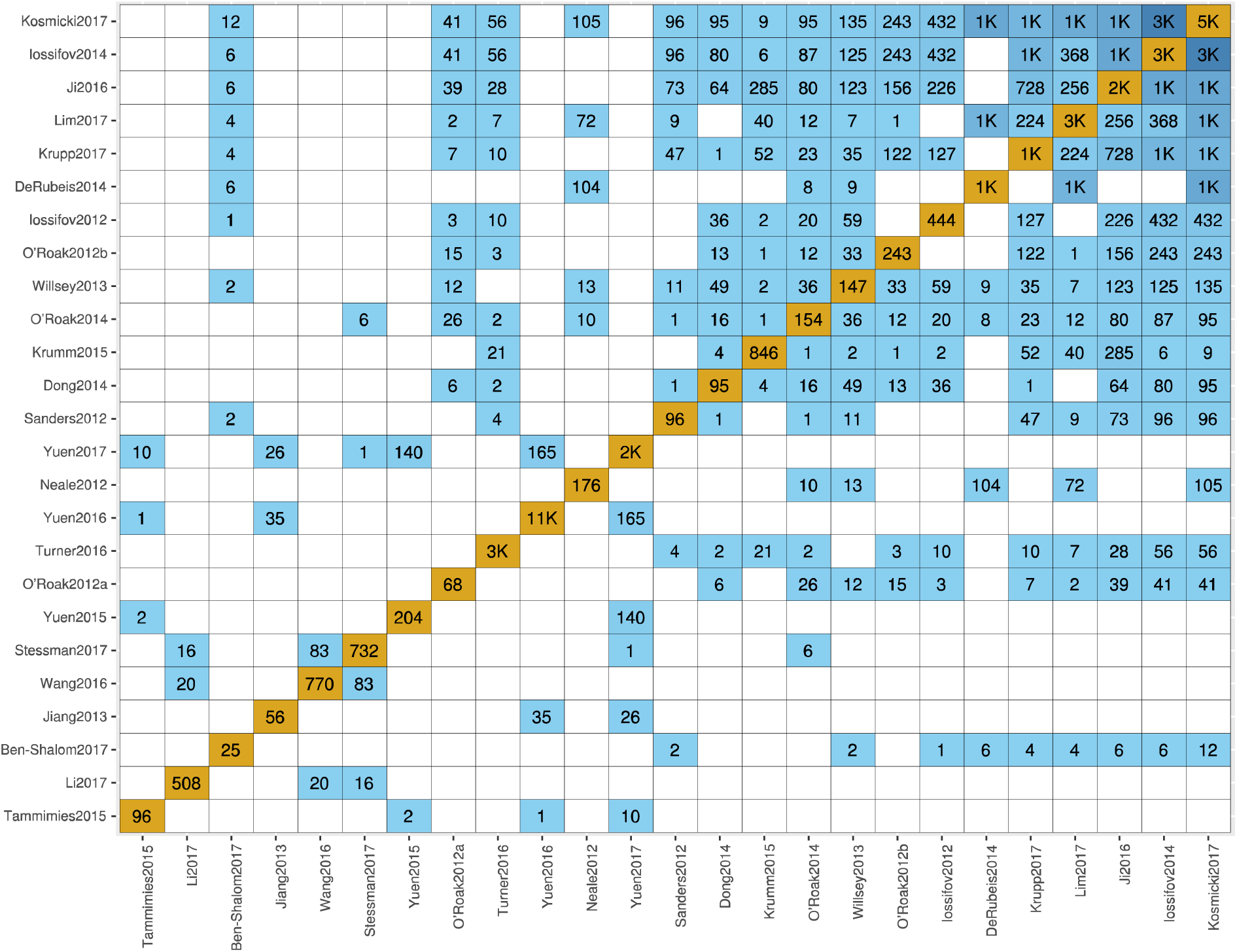
Heatmap showing the overlap between variant events reported by different papers. Cells along the diagonal represent the total number of variant events originating from each paper. Only a subset of the data is shown, for the full version see varicarta.msl.ubc.ca/stats.

Any VariCarta updates will be immediately available on the development version of the database (dev.varicarta.msl.ubc.ca), while the main site will be updated on regular bases. Previous releases of the database can be downloaded from varicarta.msl.ubc.ca/downloads. The source code for the web application and the variant processing pipeline is open source (Apache 2 license) and available on GitHub (link from varicarta.msl.ubc.ca/downloads). Questions, comments and requests should be sent to.

### VariCarta statistics

VariCarta currently contains 35,615 variant events from 8,044 subjects, collected from 50 publications. All the subjects are affected individuals (VariCarta does not collect variants from control individuals), majority of them with a reported clear ASD diagnosis and only a few with an unclear diagnosis. The latter are from publications reporting on ASD subjects among others, but without clear diagnosis label provided. Thirty-seven publications are from whole genome or whole exome sequencing studies and the remainder are from more targeted (i.e. candidate gene) studies. The variants are affecting 12,416 genes, with SCN2A being the gene with most reported variants (103 variants).

### Comparison to similar resources

We are aware of three other public databases that collect ASD variant data from scholarly literature. One of them is considered to be ASD-specific (SFARI), while the other two are more general, containing *de novo* mutations associated with multiple phenotypes (denovo-db and NPdenovo).

The SFARI gene database, a licensed version of AutDB (Basu et al., 2009) by MindSpec, is a web portal for cataloguing ASD candidate genes. The central SFARI Human Gene module contains information about human genes that have been associated with ASD, relevant references from articles, and different kinds of genetic datasets that provide evidence for linking the genes to ASD and help categorize and score genes based on the strength of that evidence. These data include genetic variants, both rare and common, reported to be associated with the disease. However, the SFARI database is not entirely ASD-specific since many of the reported variants were found in subjects not diagnosed with ASD, but other disorders, such as intellectual disability, epilepsy and general developmental delay. Since the SFARI variant table does not show the disease association, the access to ASD-specific variants is non-trivial. The variant annotation in SFARI also lacks transcript information and genomic coordinates, which makes it difficult to identify the exact genomic location of the variant and its predicted functional effect. Another concern is that, due to the multiple WES/WGS publications on the same cohorts of ASD individuals, some variants are reported more than once potentially leading to the inflation of the ASD-association evidence. Finally, the variant information in SFARI is not available for bulk download and thus cannot be used for any kind of high-throughput analysis.

denovo-db (Turner et al., 2017b) is another database of germline *de novo* variants, not limited to neuropsychiatric disorders. The latest release, from August 2018, contains data from 16 ASD studies (compared to 50 studies in VariCarta). The database provides variants’ comprehensive position annotation, damage scores, population frequency and validation status. Turner et al. (Turner et al., 2017b) discuss the variant over-reporting problem in the literature, especially in the case of heavily studied SSC cohort for which the sequencing information has been reported in multiple publications. Their solution to the problem is to avoid duplication of samples in their database, thus when a new study is imported, any samples that are pre-existing in the database would be ignored.

NPdenovo (Neuropsychiatric Disorder De novo Mutations Database) (Li et al., 2016) collects *de novo* mutations found in subjects with neuropsychiatric disorders and their sibling controls. The database provides functional annotation of the variants, information about associated genes, their brain expression patterns and co-expression. According to the website, NPdenovo was most recently updated in 2015, although their publication table (http://www.wzgenomics.cn/NPdenovo/home_detail.php) includes a few papers published in more recent years. The table lists 13 ASD-related studies, however for three of these some or all variant data were not included in the database due to the cohort overlap (as explained in the footnote of the table).

## Discussion

Here we present VariCarta, a database of literature-derived genomic variants found in ASD subjects. The motivation for developing VariCarta arose from our own research interests and inability to find a comprehensive collection of published, reconciled and well-annotated ASD-associated variants. Among the main features of VariCarta are the careful curation of variants combined with the added robustness of a data import pipeline into a relational database, with the goals of rectifying errors, standardizing reporting format, harmonizing cohort overlap, appending comprehensive annotation and tracking provenance to the original report.

Even though the prevalence of ASD is fairly high (Developmental Disabilities Monitoring Network Surveillance Year 2010 Principal Investigators and Centers for Disease Control and Prevention (CDC), 2014) the number and size of cohorts with genetic samples available for sequencing is still limited. As a consequence, some cohorts have been extensively studied. The most prominent example is the SSC cohort of 2600 families (Fischbach and Lord, 2010), which has subjects that have been reported on in at least 19 studies currently included in VariCarta. Since it is not always clear what cohorts were used in a study and there are inconsistencies in subject ID and variant reporting formats across studies, the overlap between studies’ results is not always obvious. This can lead to double-counting of variants and inflation of gene-based evidence for ASD association. Having the precise counts of reported variants per gene is important, especially considering that most ASD candidate genes have been established based on the observation of clusters of damaging mutations in multiple unrelated individuals.

One way to address this problem, as adopted by denovo-db and NPdenovo, is to avoid studies reporting on a cohort that has been used in another already curated study. While this approach may work for the obvious cases of cohort overlaps, there are many partial and more complex overlaps between studies, as illustrated in Figure 1, and consolidating these cases requires more systematic approach. In addition, studies that report on the same subjects, do not necessarily report the same variants, either due to differences in variant processing (e.g. applying different filters) or heterogeneity of variant reporting formats. Simply excluding variants from one of the studies can result in the omission of valuable information. Finally, linking variants to all the papers that report them is important for documenting data provenance and accessing additional information provided in the publication.

VariCarta puts special emphasis on harmonizing variants derived from the same cohorts of subjects by carefully linking the variants together, while still retaining the original publication and annotation data. Currently VariCarta contains 6,057 variants that have been reported more than once; some of them has been published in as many as nine papers (varicarta.msl.ubc.ca/variant?chr=20&start=49510027&stop=49510027).

Although we made a substantial effort to convert all variants into the same format sometimes the differences cannot be completely resolved; the example in Table 1 is one of the simpler cases. In these cases, we still link them together as complex events, indicating that they have been reported as overlapping variants in the same individual, and are likely to represent the same event. There are currently 327 complex events in VariCarta.

## Conclusions

To the best of our knowledge, VariCarta is the largest collection of systematically curated, harmonized and annotated literature-derived variants that is specific to ASD. Although variants can go through multiple transformation steps as they are being converted to the VariCarta uniform format, these steps are documented and the original variant information is retained and readily accessible. Our curators are always on the lookout for new relevant publications, which are being continuously added to the database. While whole genome and whole exome studies are of primary interest, we will continue to add gene-targeted studies that are aligned with our research interests or requested by the research community. We hope that the ASD research community as well as clinicians working with ASD subjects will find VariCarta to be a useful resource.

## Authors’ contributions

MB implemented the data processing pipeline and contributed to software development. MJ designed and engineered the web application and relational database model. NH participated in the initial data collection. JP carried out curation of all publications and performed data processing tasks. SR and PP conceived the study. PP provided overall project leadership. SR provided project management and led database design and testing. SR drafted the manuscript with input from PP and MB. All authors read and approved the final manuscript.

## Acknowledgements

We would like to thank the authors of publications included in VariCarta who communicated with us to provide clarification or additional information.

